# DBSOMA: A Machine Learning Method that Identifies Chemical Modulators of Transcriptional States Uncovers Effectors of Beta-Cell Maturation

**DOI:** 10.64898/2026.01.30.701818

**Authors:** Timothy R Kunz, José Rivera-Feliciano

**Affiliations:** Department of Stem Cell and Regenerative Biology and Harvard Stem Cell Institute (HSCI) Harvard University, Cambridge, MA 02138, USA

**Keywords:** Machine Learning, Self-organizing Map (SOM), Density Based Clustering of applications with noise (DBSCAN), in silico chemical screening, gene expression patterns, protocol generation, beta cell maturation, Stem cell derived islets (SC-Islets), in silico chemical screening, gene expression patterns, islet function

## Abstract

The effects of perturbation on a biological system can be readily measured in terms of transcriptional changes. However, despite a wealth of transcriptional perturbation response data, there are currently few methods to draw equivalence between the many biological systems used to generate that data and a specific system of interest. Here we use density analysis of transcriptional correlations to computationally predict whether a given perturbation readout is relevant to Stem Cell derived islet (SC-Islet) maturation. The approach, Density Based Self-Organizing Map Analysis (DBSOMA), first learns patterns of gene expression represented in scRNA-seq sets by clustering genes with the Self-Organizing-Map (SOM) algorithm. Perturbation expression profiles and other gene lists are then projected onto the SOM grid, where the degree of clustering is determined by the Density-Based Spatial Clustering of Applications with Noise (DBSCAN) algorithm. We applied DBSOMA to SC-Islet maturation and identified known and novel regulators of β-cell maturation. This workflow can be applied broadly to biological systems where single-cell RNA-sequencing data is available, and a desired outcome can be represented in transcriptional changes.

## Introduction

In-vitro differentiation of human embryonic or induced pluripotent stem cells into SC-Islets is a promising avenue as a replacement therapy for the treatment of type I diabetes. SC-Islets are produced through a well characterized protocol, wherein stem cells are differentiated through stages that mimic embryonic development with specific cocktails of chemical and growth factor treatments [1]. The resultant SC-Islet organoids are composed of pancreatic endocrine cells, such as insulin producing β-cells and glucagon producing α-cells and enterochromaffin cells. [1, 2]. The in-vitro derived islets have been characterized to have a fetal-like signature and to be less responsive to glucose than primary human islets [3, 4]. However, upon transplantation of islet organoids, the cells rapidly acquire functional maturity [5, 6]. While this suggests that maturing the cells *in vitro* is not necessary for their use as a cell replacement therapy, we can still gain insights on the maturation process by studying the SC-Islet cultures. Additionally, effectors of functional maturation i*n vitro* may represent promising candidates for improving beta cell function in type II diabetic contexts, where beta cell function is impaired [7]. Several efforts have been made to mimic this maturation process *in vitro*, and certain protocols report successful improvement of functionality through an additional stage of differentiation with compounds to induce maturation signatures [6, 8, 9].

We aimed to gain a greater understanding of the maturation process and to expand the repertoire of maturation-inducing chemicals that can be added to the protocol by consulting the dense perturbation response data from the CMAP L1000 project [10]. The project involved perturbing multiple different cell lines with over 25,000 agents, including 19,000 chemical perturbagens. The Ma’ayan lab’s SigCom LINCS resource has conveniently distilled the L1000 response data into lists of the 250 most up and down regulated genes for every perturbation scheme tested, resulting in ∼1.4 million transcriptional response signatures from chemical perturbagens alone. Additionally, the resource includes sets from other perturbations, such as CRISPR knockout, shRNA, siRNA, antibody treatment, overexpression and ligand treatments [11].

Querying these datasets with this gene up/down datatype is challenging. Most searches are done based on co-occurrence of genes by name. Analysis considering gene similarity, rather than by name, with the use of deep learning, has been shown to outperform analysis based on identity alone [12]. The perturbations themselves also introduce caveats, since silenced genes may affect biological processes outside of their annotated role, and pharmacological perturbations may have promiscuous targets, leading to unexpected responses [13-15].

There is also the challenge of determining which datasets are apt for use in a system of interest. For example, it may be wise to select data collected from cancer lines derived from a primary tissue of interest, but still the cell lines may behave in unexpected ways in response to perturbation. Still, we hypothesized that, broadly, certain cell lines will respond to perturbation in ways analogous to SC-islets. We therefore used single cell sequencing data from SC-Islets to contextualize perturbation responses that mimic gene regulatory networks at play in the β-cell maturation process.

Single cell RNA sequencing analysis is traditionally done in pipelines with dimensionality-reduction algorithms, like TSNe and UMAP, and focuses on characterizing cellular heterogeneity [16]. Alternatively, approaches such as non-negative matrix factorization (NMF) and independent component analysis (ICA) can cluster genes with similar expression patterns and similar roles [17, 18]. Another approach for analysis, the Self-Organizing Map (SOM) algorithm, has been used to analyze large micro array datasets to cluster genes with similar expression patterns [19]. For our purpose, the SOM offers an important advantage that training is restrained to a rigid grid, which is apt for the application of the density calling Density-Based Spatial Clustering of Applications with Noise (DBSCAN) algorithm. DBSCAN has been successfully applied to single-cell data analysis to identify clusters of like cells [20], but it is more commonly applied to spatial datatypes [21, 22]. This spatial analysis allows for the identification of similar patterns in the gene correlation high dimensional space without a dependency on gene sets containing exact gene matches. In this work, we establish a framework to identify regions on the SOM of common pattern changes between perturbation responses and desired outcomes, without reliance on the co-occurrence of impacted genes, to predict novel chemical inducers of β-cell maturation transcriptional programs.

## Results

### DBSOMA: algorithm overview

First, single-cell RNA-sequencing data from SC-Islet culture is used to train a self-organizing map. The resultant map represents a latent space in which genes with similar expression patterns in the initial cell culture are grouped together (Figure 1A). Genes are assigned to individual nodes in a toroidal grid of hexagonal nodes.

**Figure 1:**
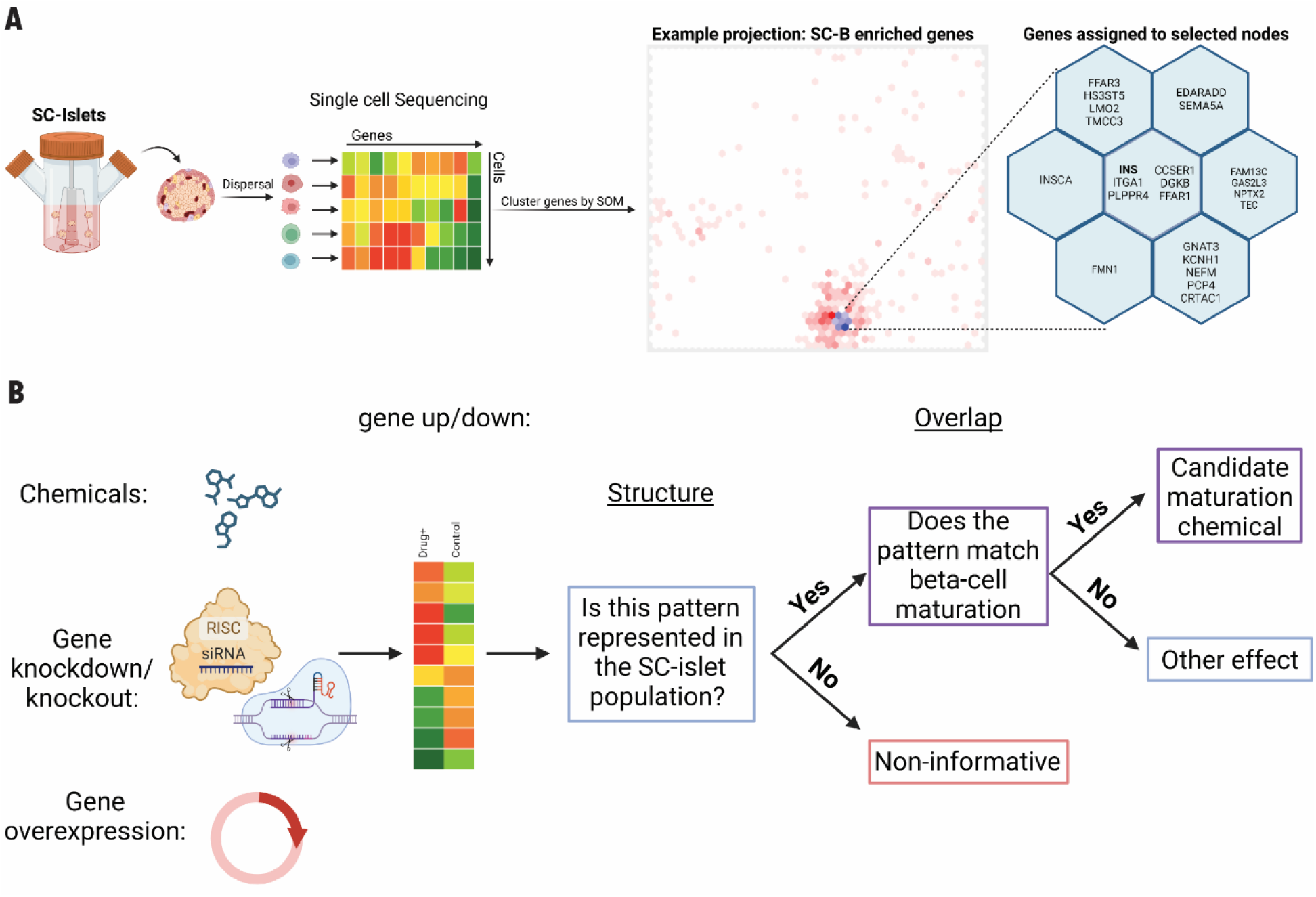
DBSOMA: Algorithm overview. Single-cell sequencing from a cellular population of interest is used to generate a gene x gene pairwise-correlation matrix, which is then used as input to train a Self-Organizing Map (SOM), clustering genes onto a grid according to their similarity in the pair-wise correlation data space, where a given list of genes can be projected to see where on the map they were assigned. An example of SC-β enriched genes is shown (A). This SOM output allows us to automate a process where perturbation transcriptional responses are scored according to their ability to form structure on the Self-Organizing Map and whether that structure overlaps with a desired outcome (B).

Next, the self-organizing map is used to define a target pattern of gene expression; in our case, this is β-cell maturation. L1000 perturbation transcriptional responses are then projected onto the SOM. First, we ask whether any pattern is represented on the SOM, and if so, does the pattern overlap with the target pattern of maturation genes (Figure 1B).

### Clustering of genes from single-cell RNA-seq of *in vitro* differentiated islets reveals SOM region associated with increased maturation

A SOM was trained on single cell RNA-sequencing data produced by Veres et al. from stage 6 SC-Islets [2]. This SOM is referred to as the SC-Islet SOM. Specific lists of genes were then input to visualize where those genes reside on the map and whether they form a cluster. As controls, we projected sets of genes associated with well-established cellular processes and of known cellular identity, including cilia formation, ribosomal proteins, cell cycle regulation, and fetal pancreas identity, onto the SOM. As expected, we produced patterns (Supplemental Figure 1). Therefore, we confirmed that genes involved in different processes result in differential spatial patterns represented in the output space, and we hypothesized that some of these patterns may be reproduced by drug induced transcriptional changes.

We next sought to define a pattern representative of β-cell maturation. Augsornworawat and colleagues produced upregulated gene lists for many cell types in islet clusters, relative to the other cell types, from various islet sources [23], including pre-and-post transplantation SC-β, as well as primary human β-cells. Each of the beta cell enriched gene lists from the three islet sources were projected onto the SC-Islet SOM (Figure 2A). From all three datasets, we see a prominent cluster which includes key β-cell genes like insulin, glut2 and IAPP. However, post-transplantation SC-Islets and primary human islets also have a second prominent cluster that contains several genes known to be upregulated as β cells mature, including SIX2, GIPR and GLP1R [24, 25]. Post-transplantation SC-β also form a third cluster containing many ribosomal protein genes. We observe that the cluster containing maturation genes appears in the more mature cell sources, namely post-transplantation SC-β and primary human β-cells, so the region of the map defined by this cluster represents a target for perturbation responses which may induce maturation. We refer to the set of genes defining this cluster as the maturation neighborhood (extended data table 1). As an alternative approach, we mapped the lists of genes differentially expressed between primary β-cells and SC-β (Supplemental figure 2). We observe the lists of upregulated genes to form a structure apparently similar to the cluster of ribosomal protein genes in Sup. Figure 1.

**Figure 2:**
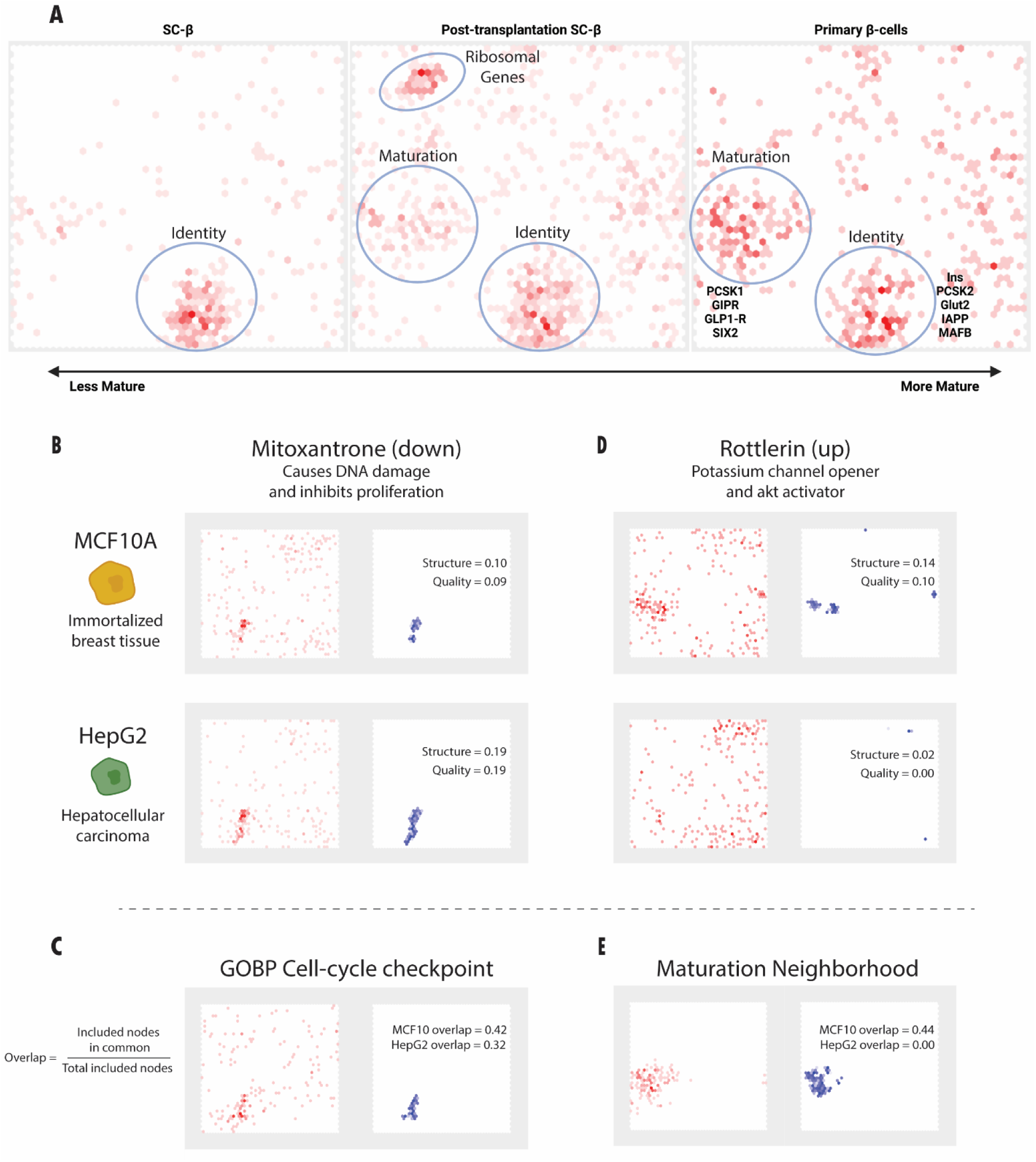
Mapping and scoring perturbagen responses to a maturation neighborhood. Projection of upregulated genes in less mature SC-Islets, more mature post-transplantation SC-Islets and most mature primary human islets also form apparent clusters on the SOM, with a structure containing known maturation genes becoming more apparent in more mature islet sources (A). This cluster defines our maturation neighborhood target. Applying DBSOM to perturbation response gene list projections allows us to determine the relative structure and quality of the data on our map for different drug-cell line combinations (B, D). The effect of the drug can then be estimated by the degree of overlap with comparison gene-lists (C, E). Input projections are shown in the left panels in red and DBSCAN *included* nodes are displayed in right panels in blue (B-E).

Downregulated genes and the combined gene lists produced considerably more disperse mapping on the SC-Islet SOM compared to our defined maturation neighborhood (Supplemental figure 2). The post-transplantation β-cells were also observed to have a cluster of upregulated ribosomal protein genes (Figure 2A, middle panel). While this upregulation of ribosomal protein genes in the more mature β-cell sources may be interesting, we decided to focus our efforts on targeting the identified maturation neighborhood.

### DBSOMA allows for the scanning of perturbation data sets for overlapping density with the maturation neighborhood on the SOM

We hypothesized that finding perturbations which produce regions of co-density on the SOM with the maturation neighborhood would indicate compounds capable of inducing maturation transcriptional programs. To find regions of density, we applied DBSCAN to perturbation response sets. The algorithm iterates through a list of genes searching a radius around the node to which each gene is assigned and *includes* nodes that are found within a minimum number of checked radii. Nodes above that threshold are *included* (Supplemental Figure 3 A&B). The number of included nodes is then used to calculate two metrics describing the density. First, the *structure* metric is calculated as the ratio of *included* nodes to number of input genes, then the *quality* of that structure is determined as the ratio of the number of genes assigned to *included* nodes to the number of input genes (Supplemental Figure 3A&B). Higher *structure* and *quality* values indicate a relatively higher degree of clustering of the input list of genes on the SOM with less noise, indicating that this pattern of gene changes is reflected in the initial SC-Islet RNA-seq dataset. However, defining a high or low value for these metrics depends on the context of the input data sets and the SOM on which sets are projected.

Next, we examined sample gene up/down lists taken from SigCom Lincs1000 data set [11] and subjected them to density calling by DBSCAN on the SC-Islet SOM to see whether any produced regions of density and if so, whether any had the expected regions of co-density with the maturation neighborhood. The lists of down-regulated genes from treatment with the general DNA damaging and proliferation inhibitor mitoxantrone formed an apparent cluster in both MCF10A (mammary cell line) cells and HepG2 (human hepatocellular carcinoma) cells, with the HepG2 set scoring higher in structure and quality (Figure 2B). Predictably, this region of density overlaps highly with a projection of cell-cycle checkpoint related genes (Figure 2C). Based on these results, we would correctly predict that mitoxantrone inhibits the cell cycle. Treatment with rottlerin, a mitochondrial uncoupler and membrane depolarizer which inhibits PKC-δ [26, 27], enriched genes that formed a cluster in MCF10A cells, but not HepG2 cells, which was reflected visually and by the lower *structure* and *quality* scores in the HepG2 (Figure 2D). The region of density formed by the list of genes upregulated in MCF10A in response to rottlerin overlaps with the maturation neighborhood (Figure 2E). We would therefore predict that rottlerin treatment will increase the maturation transcriptional program in SC-β. We were reassured by this example of our desired outcome: a perturbation response from a cell line that forms a cluster on the SC-islet SOM with relatively high *structure* and *quality* that overlaps with the maturation neighborhood. DBSOMA therefore allows for the quantification of the degree of clustering a given gene list produces with a *structure* and *quality* metric, as well as a measure of the amount of overlap the resultant density has with the desired maturation neighborhood. Importantly, this process does not rely on co-occurrence of genes between target and perturbation gene lists and allows for a high-throughput, automated search of gene lists based on these metrics.

### Identification of maturation inducing agents from the L1000 chemical perturbations dataset

The Ma’ayan lab’s SigCom LINCS provides nearly 1.4 million up or down regulated gene signatures from chemical perturbation data distilled from the L1000 project [11]. Signatures were first evaluated for their degree of *structure* and *quality* and then for their overlap with the β-cell signature (Figure 3A). The *structure* (Figure 3 B&D) and *quality* (Figure 3 B&E) values were calculated as described previously. To minimize the occurrence of false positives, we also generated 100,000 lists of genes randomly selected from the SigCom input lists. These randomized lists were projected onto the SOM and evaluated for our clustering metrics (Supplemental Figure 4). Thresholds of 0.075 for *structure* and 0.57 for *quality* were set as roughly double the highest scores from the randomized gene lists, ensuring the false discovery rate is significantly lower than 1 in 100,000. Perturbation response signatures passing those thresholds were then evaluated for their degree of *overlap* (Figure 3 C&F) with the maturation neighborhood. The threshold for significant overlap of 0.1 was determined ad hoc, based on manual evaluation of edge case overlaps, and yielded 224 putative maturation-inducing signatures. Signatures could then be ranked by the *product* of the *structure*, *quality*, and *overlap,* to identify those with relatively high scores in all metrics (Figure 4G). Overall, from the initial 1.4 million chemical response signatures, 21 thousand passed structure and quality thresholds, and of those, 224 were found to be overlapping with the maturation neighborhood. Of the 290 cells lines with at least 1 associated signature, only 38 produced a candidate maturation neighborhood. Among the individual cell lines, several were found to be significantly overrepresented by hypergeometric testing (Figure 4H). Interestingly several mammary epithelial lines, including MCF10A, MDAMB231 and T47D, which may suggest this cell type is generally more capable of transcriptional changes analogous to beta cell maturation. Of the 224 chemical signatures predicted to induce maturation, there were 201 unique drugs with 23 duplicates from either multiple cell lines or multiple dosages. The complete list of hit signatures is included (extended data table 2).

**Figure 3:**
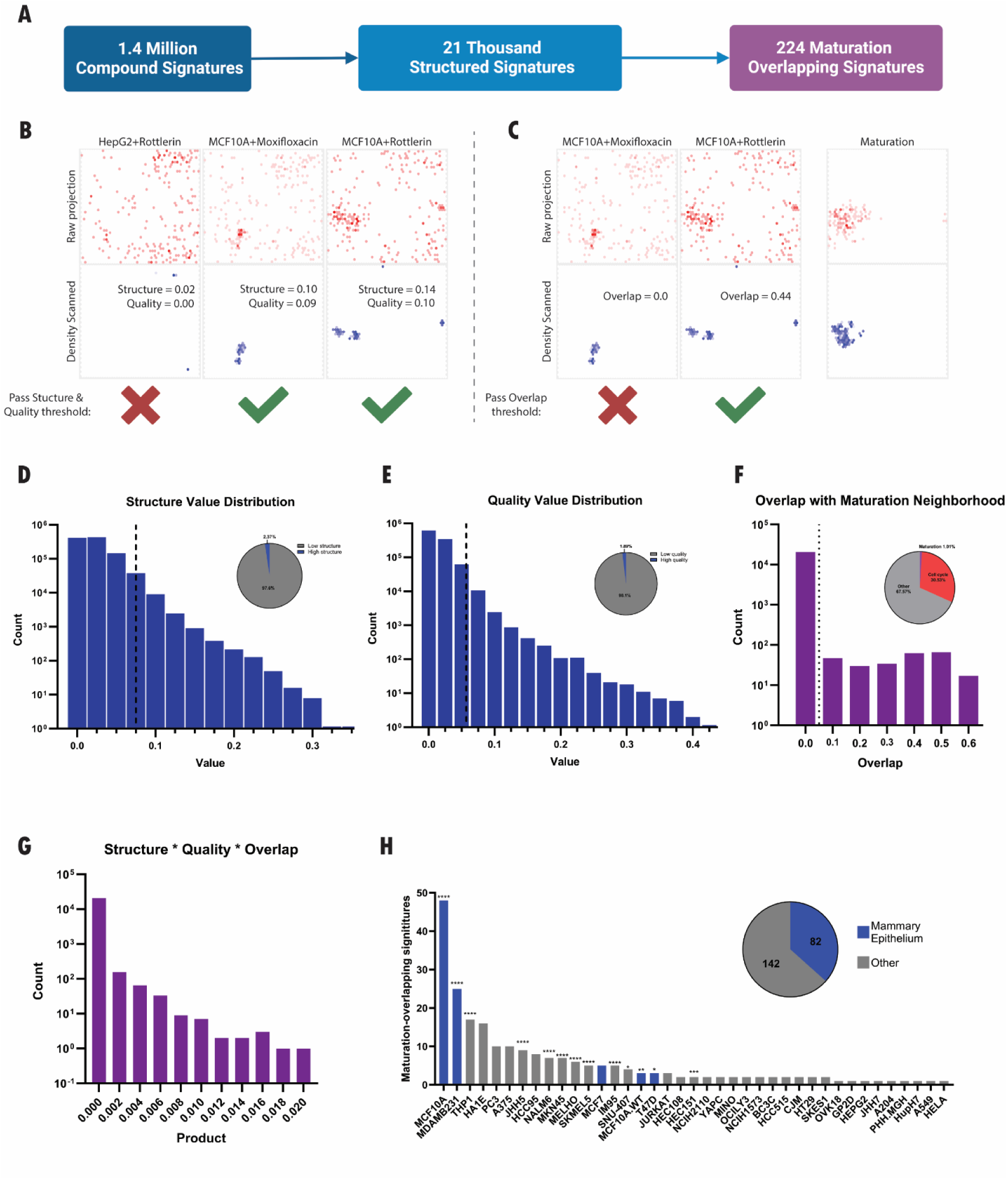
In silico screening of L1000 perturbagen signature yields candidate maturation signatures. L1000 gene response signatures were subjected to a 2-step screening process, first for those forming structures on the SOM then for those structured signatures that overlap with the maturation neighborhood (A). Previous example projections and their relevant values for screening are shown in B & C. First, signatures below thresholds (dashed lines) for *structure* (D) and *quality* (E) were eliminated. Signatures were then assessed for their degree of *overlap* with Beta Cell maturation neighborhood (F). Multiplying *structure*, *quality* and *overlap* metrics can be used to calculate an overall score for each signature (G). The 224 signatures passing thresholds were produced largely by mammary epithelium cell lines; significant overrepresentation was assessed by Benjamini-Hochberg hypergeometric testing against the total number of signatures in the L1000 dataset for each line (*= p<0.05, **= p<0.01, ***= p<0.001, ****=p<0.0001) (H).

**Figure 4:**
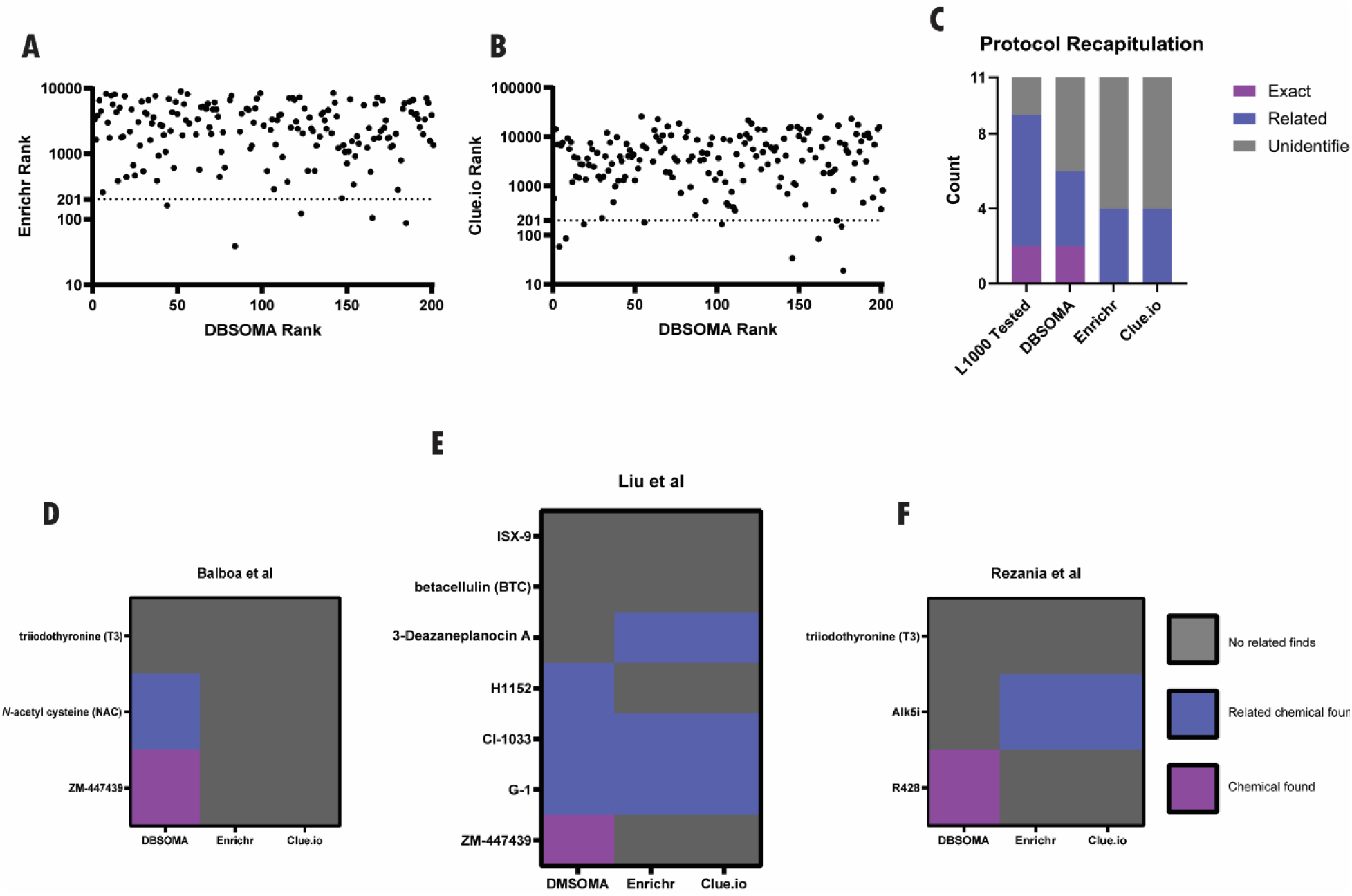
DBSOMA compared to similar methodologies. The 201 putative maturation-inducing chemicals predicted by DBSOMA were ranked by Enrichr (A) and Clue.io query app (B). Next, the top 201 ranked compounds by each method were compared to the 11 unique compounds used in the three in-vitro maturation protocols and compared to the compounds directly tested in the L1000 dataset (C). Each protocol is individually examined as well (D-F). Compound finding is scored based on ability to find the exact compound, a different compound with a similar annotated function (related), or finding no related compounds.

### DBSOMA identifies compounds not predicted by comparable methods, including known maturation compounds

To evaluate the predictions generated by DBSOMA, we compared the approach to the Clue.io query app and the Enrichr tool. These tools utilize the same L1000 datasets and were given the same initial target list of primary islet upregulated genes used to identify the maturation cluster in Figure 2. First, we evaluated the rank of each of the 201 unique maturation chemicals according to Enrichr (Figure 4A), and the query app (Figure 4B). In both cases, DBSOMA-identified compounds are generally ranked lower in the comparison methods, and few compounds are highly ranked by both DBSOMA and a comparison method. Therefore, the DBSOMA methodology identifies compounds that would not be indicated by these other methods.

Next, we compared the lists of predicted chemicals from the various methods to compounds used in three published protocols for the induction of *in vitro* β-cell maturation, Balboa et al. [6], Rezania et al. [8], and Liu et al. [9], to determine the ability of each approach to identify known maturation-inducing compounds (Supplemental table 1). Of the 11 unique compounds utilized in these protocols, only two, the aurora kinase inhibitor ZM447439 and the Axl inhibitor R428, were directly tested on cell lines in the L1000 project. For several of the protocol compounds that were not included in the L1000 dataset and therefore could not be directly identified by the various approaches, other compounds with related functions were identified. We determined whether DBSOMA, the Clue.io query app and Enrichr predicted within the top 201 scored chemical perturbagens the exact compounds, a compound with a similar function, or no related compounds for each chemical from the protocols. DBSOMA outperformed both other methods, identifying more exact and related chemicals used in these maturation protocols (Figure 4C). Additionally, only DBSOMA was able to identify ZM447439 (Figure 4 D-E) and R428 (Figure 4F). Together, these results show that DBSOMA produces a set of predicted perturbations different from related methodologies and, importantly, it was able to predict perturbations known to induce β-cell maturation.

### Maturation overlapping signatures from chemical and genetic perturbations affect pathways and processes associated with beta cell maturation

We next performed a literature review of the compounds indicated as potentially maturation inducing by DBSOMA but not associated with known maturation protocols. We determined that many of the compounds, including ciclopirox, epirizole, flutamide, gemifloxacin, imatinib, LY-294002, milnacipran, MK-2206, moxifloxacin, neratinib, rimonabant and roscovitine, had been previously implicated in islet function, endocrine differentiation and glucose homeostasis (Supplementary Table 2).

In addition to chemical perturbations, we applied the DBSOMA workflow to the SigCom collection of genetic perturbations by CRISPR knockout and overexpression. This approach identified 35 of the 281 thousand knockout signatures and 17 of the 68 thousand overexpression signatures as overlapping with the maturation neighborhood (Supplemental Figure 5A). These lists of gene targets were combined and subjected to GO:term enrichment analysis, revealing enrichment of several maturation-relevant GO:terms, including regulation of endocrine pancreas development, and circadian rhythm (Supplemental Figure 5B) [28]. Together, these results further suggest that DBSOMA identifies known regulators of β-cell function and maturation.

### Drug treatments have predicted effects on maturation gene mRNA expression

Because the DBSOMA approach was able to correctly identify chemical and genetic perturbations known to induce *in vitro* maturation of β-cells, we hypothesized that the other identified perturbations would induce an *in vitro* maturation transcriptional signature. To test this, we differentiated 3D cultures of SC-Islets until the stage 5 endocrine progenitor stage. Clusters were then dissociated and plated into 2D cultures, and the protocol was continued with the standard differentiation cocktail with the addition of a DBSOMA-identified compound or a control volume of Vehicle (Figure 5A). Differentiation continued through the end of stage 5 and for 7 days of stage 6. We tested only a subset of 7 of the 201 predicted compounds, which we selected to encompass a range of annotated mechanisms of action.

**Figure 5:**
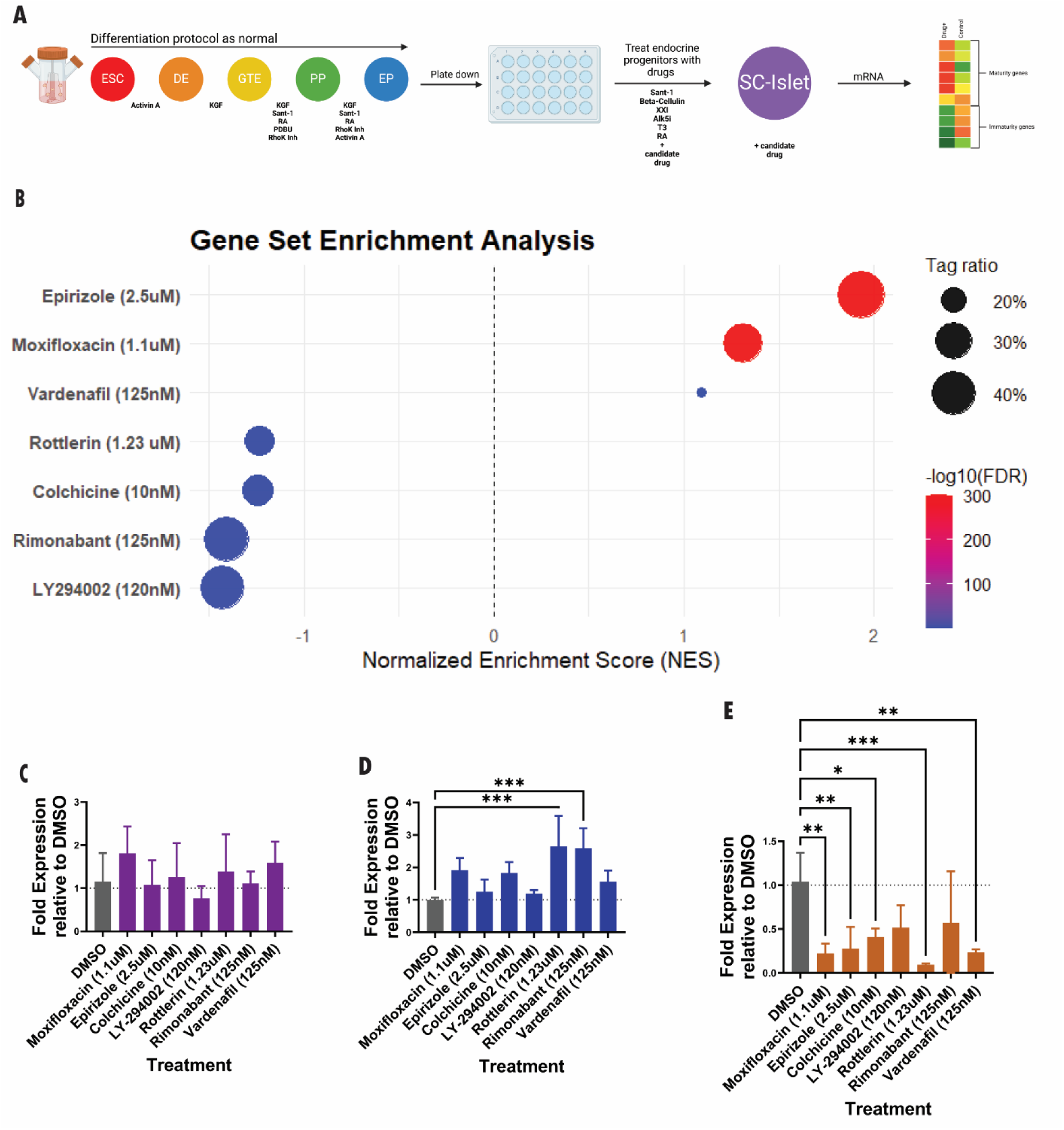
Screening for maturation inducing compounds among identified candidates. SC-B cells were grown in 3D culture until the completion of stage 4, then cells were dispersed and replated. Stage 5 cells were then treated with standard cocktail plus a candidate compound, through the completion of stage 5 and for the 7 days of stage 6 (A). Bulk RNA-seq was performed on whole islet cultures and differential gene expression of genes comprising our maturation neighborhood was examined for each drug treatment relative to DMSO control by gene set enrichment analysis (B). Expression for specific genes was examined by qPCR for changes in expression of insulin (C), MAFA (D), and NPY (E).

We then performed bulk RNA sequencing on the resultant cultures, which included other cell types in addition to SC-β. To evaluate whether drugs were upregulating maturation neighborhood transcripts, we performed gene set enrichment analysis for each treatment relative to Vehicle treated controls (Figure 5B). We observe that moxifloxacin and epirizole statistically significantly enrich for upregulation of maturation neighborhood genes.

We also specifically measured expression by qPCR of insulin (Figure 5C), the well characterized maturation marker MAFA (Figure 5D) [5, 29] and the immaturity marker neuropeptide Y (Figure 5E) [30]. While treatments did not significantly change insulin expression, treatments with rottlerin and rimonabant significantly increased MAFA expression. Conversely, many of the treatments decreased NPY expression. These patterns of increased maturation neighborhood genes, increased MAFA expression and decreased NPY expression suggest that the drugs are inducing a β-cell maturation transcriptional program in beta cells.

Finally, we analyzed the top 500 differentially upregulated genes relative to Vehicle for each treatment and projected these gene lists onto the initial SOM (Figure 6A), and calculated *structure* (Figure 6B)*, quality* (Figure 6B), *overlap* (Figure 6C) and *product* (Figure 6D). We observe both visually, and based on our analysis metrics, that genes upregulated by epirizole and moxifloxacin treatments produce clusters on the map significantly overlapping with the maturation neighborhood. Because sequencing was performed on whole cultures of treated cells, drugs not producing this pattern may still be inducing maturation in β-cells, but the signal may be hidden by stronger effects in other cell types. Overall, these results suggest that compounds predicted by DBSOMA can promote the upregulation of maturation genes in SC-β.

**Figure 6:**
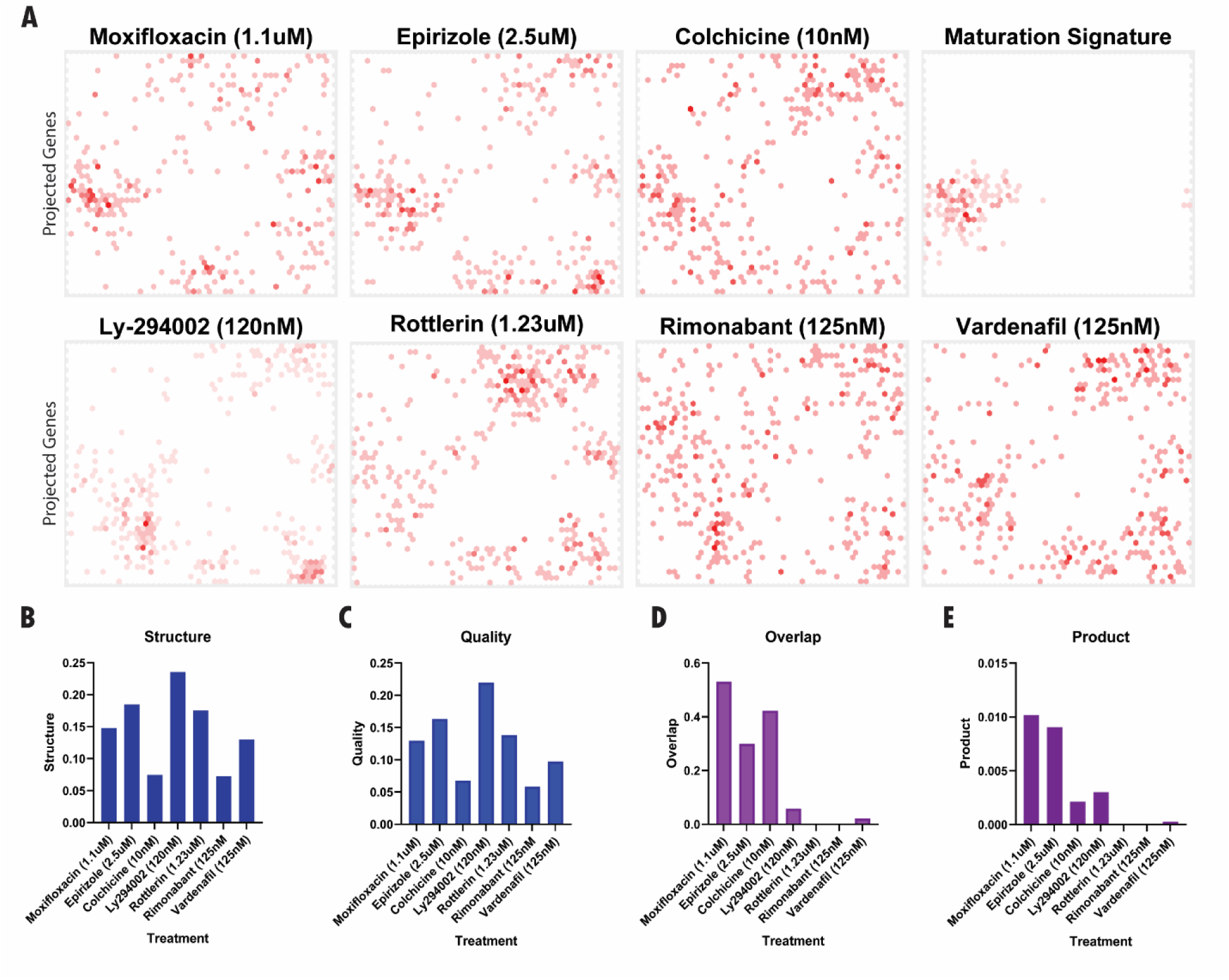
SOM analysis of chemically induced transcriptional changes in SC-Islet cultures. The top 500 upregulated genes for each drug treatment, relative to DMSO treated controls, were projected onto the trained SOM and compared to the maturation neighborhood projection (A). Each projection was then evaluated for structure (B) and quality (C), then overlap with the maturation neighborhood (D) and product of those values (E) were calculated.

## Discussion

We established a two-step framework for contextualizing perturbation transcriptional output data from disparate cell types to stem-cell derived β-cells. We used it to identify known and novel regulators of a gene cluster associated with increased beta cell maturation. Among the drugs tested in vitro on SC-islets, several showed signs of increased transcriptional maturation, including increased MAFA expression, decreased NPY expression and a general upregulation of our identified maturation neighborhood genes. This analysis was performed on bulk RNA from SC-islets, including beta cells and other cell types, so future work involving single cell sequencing will be required to address what transcriptional changes are specific to the beta cells. Of the 201 DBSOMA identified compounds, only 7 were tested here. Other compounds from this set may have more profound or specific effects on the beta cells. It also remains to be established whether these compounds can affect functional maturation *in vitro* or *in vivo*, which will require further experimentation. Overall, though, we were able to make predictions on transcriptional outcomes for the SC-islet system based on SC-islet single cell sequencing and the L1000 gene list data set.

This approach is flexible to any system in which scRNA-seq data is available and a robust set of gene targets can be defined. In cases where the set of gene targets is not previously defined, analysis of where related differentially expressed genes cluster on the SOM can be used to define a target, as we did to define the maturation neighborhood. With the massive expansion of single-cell data repositories, there are ever increasing use cases for this workflow, including differentiation to other cell types, cancer evasion, aging, metabolism or disease signatures.

In both benchmarking and predicting novel maturation compounds, this study was limited to the compounds tested in the source L1000 data. For example, only 2 of the 11 compounds used in reference maturation protocols were directly detected. However, the workflow is not limited to L1000 data, and indeed there are numerous other sources of gene up/down perturbation response data that could be used in addition to the L1000 data for future applications.

We observed that mammary epithelium cell lines were overrepresented among cell lines generating a maturation neighborhood overlapping signature. This may suggest that using these mammary epithelium lines could be used for screening additional compounds prior to use in the complicated SC-islet system. More broadly, the approach could identify proxy cell lines for expanded screening on more easily manipulated cell lines for whatever system is being examined.

The SOM algorithm was selected due to the ease with which DBSCAN could be implemented on the output. However, this approach could be modified to use other dimensionality reduction algorithms, other density scanning approaches, or both. Additionally, we hypothesize that density analysis of the high dimensional cell x gene count matrix of single cell RNA-seq data could be used effectively. However, for this application, the visual element of the SOM allowed for by-eye target cluster identification, which may be more difficult in high dimensional space.

Overall, this analysis establishes a framework for leveraging scRNA-seq data to contextualize perturbation results to a specific system of interest and offers a broad set of use cases.

## Methods

### Code availability

The open-source Java implementation used for pairwise correlation matrix generation, SOM training and DBSOMA implementation can be found at https://github.com/trk5150/DBSOMA. This repository also includes several python scripts used for data processing.

### scRNA-seq Data

Single cell sequencing data for stage 6 stem cell derived islets, were sourced from Veres et al (GSE114412) [2]. Genes with counts lower than 0.5% of average read count were removed from analysis. A pairwise Pearson correlation matrix was then generated for all remaining genes.

### Self-Organizing Map

Our SOM implementation, modified from Kunz et al, 2021 [31], defines the output space on a toroidal grid with hexagonal nodes, each having 6 neighbors. This output space can be visualized as a 2D grid, with connected borders and corners. Each node contains an N-dimension weight vector, where N is the number of genes from the input gene x gene correlation matrix. These vectors are initialized to be equal to a randomly chosen data point.

The batch SOM training process involves two-step iterations of an assignment step and an update step. During the assignment step, each gene’s data vector is compared to each node’s weight vector, and is assigned to the most similar node, as measured by Pearson correlation. Node weight vectors are then updated according to the formula:

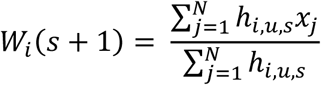

Where: s is current iteration; *W_i_* is the weight vector of node *i*; *N* is the number of datapoints (genes) in the training set; *x_j_* is the data vector for data point *j*. and *h_i,u,s_* is the neighborhood function defined as follows:

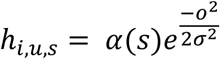

Where: *u* is the SOM node to which datapoint *j* is assigned, *o* is the distance on the SOM grid between node *i* and node *u*; *σ* is the variance of the Gaussian kernel for the current iteration; and α(s) is the learning rate for the current iteration. The learning rate shrinks throughout training from 1.0 to 0.01, and the variance parameter shrinks linearly from 3 to 0.2. The SOMs herein were trained for 100 iterations.

SOM training requires two files: a tab-delimited pairwise correlation matrix and a line split list of genes corresponding to each row in the correlation matrix. Training outputs two files: one contains metadata describing the training and input parameters and another with the trained self-organizing map, with genes listed for each node coordinate on the output grid space.

### Density-based spatial clustering of applications with noise

The implementation of the DBSOMA algorithm can be found at the same GitHub repository referenced above. As discussed further in results, gene lists are projected onto the SOM grid, and nodes are weighted based on the number of listed genes assigned to each node. A radius is then drawn around each node, and nodes with weight within that radius above threshold are included. A radius of 1 and a minimum threshold of 5 was determined ad hoc based on by-eye evaluation of clusters to give results most representative of initial cluster shape for the grid size and gene list size used for analysis here. However, changes in either grid size or gene list size may require changing the radius and threshold parameters.

### Gene list data sourcing

Analysis of gene lists using the DBSOMA algorithm on the SOM can be done with any list of genes. Very small (<100 genes) or large lists (>1000 genes) tend to produce worse results based on our analysis. Gene lists for perturbations analyzed herein were obtained from the Maayan lab’s distillation of the L1000 perturbation signatures, found here: https://maayanlab.cloud/sigcom-lincs/#/Download. Additional gene lists were obtained from the GSEA data base (https://www.gsea-msigdb.org/gsea), and the maturation target signature was generated through analysis of the lists of most highly upregulated genes in pre- and post- transplantation SC-β and cadaveric β-cells obtained from Augsornworawat et al [23].

### Gene enrichment analysis

Gene set enrichment analysis was performed on targets identified from CRISPR knockout targets or overexpression. Biological process enrichment analysis was performed using a custom permutation-based analysis to account for the highly skewed testing distribution in the L1000 datasets. The list of predicted gene perturbations was compared to each biological process list with fewer than 1000 and greater than 5 terms, and then 100,000 lists of equal size were randomly selected from the L1000 testing background. Enrichment was calculated relative to the randomly generated gene sets.

### Candidate benchmarking

Benchmarking was performed against the Maayan lab Enrichr tool [33] and the Clue.io query app [10]. Both tools were provided with the same target gene list of upregulated in primary β-cells that was used to define the target maturation cell state in on our Self-Organizing maps. Query app accepts a maximum of 150 genes as input, so only the top 150 from the primary beta cell upregulated genes were used. Query app rankings were preprocessed before comparison to remove duplicate compound signatures, and only the highest scoring signature for each compound was considered for determining rank. A complete list of rankings by both methods for each of the 201 DBSOMA identified compounds can be found in extended data table 3. The comparison methods were also evaluated using the differentially expressed genes between primary β cells and SC-β, and each performed similarly (data not shown). Related chemicals were defined as compounds annotated to have the same target in the L1000 metadata, and these implied equivalencies can be found in extended data table 4.

### Chemical sourcing and preparation

| Compound | Vendor | Cat. No. | Lot No |
| --- | --- | --- | --- |
| Colchicine | Thermo Scientific | 22712001 | A0448324 |
| Ly-294002 | Adipogen | 154447-36-6 | A00094 |
| Moxifloxacin | TCI American | M2479 | XQHXE-SD |
| Epirizole | Toronto Research Chemicals | TRC-M225093 | 2-PVA-86-1 |
| Rottlerin | Enzo | ALX-350-075-M010 | 01172002 |
| Vardenafil | Selleck Chem | S2515 | S251502 |
| Rimonabant | Selleck Chem | S3021 | S302103 |

Compounds were resuspended in DMSO and diluted to final concentrations indicated by identified hit signatures such that 1ul of DMSO suspended compound was added to culture. In the cases of moxifloxacin and rottlerin, where the compound produced a hit signature at multiple dosages, the indicated dosages were averaged for testing.

### Cell Culture

Differentiation of human pluripotent stem-cells was carried out as previously described [1]. Pluripotent stem-cell lines were obtained from stocks maintained by Rivera-Feliciano. The insulin-red reporter line was generated as a 3’-UTR knock-in. Pluripotent stem-cells were maintained in suspension clusters using mTeSR1 Stem Cell Technologies, 85850) in 500-ml spinner flasks (Corning, VWR) spinning at 70 r.p.m. in an incubator at 37 °C, 5% CO2 and 100% humidity. Cells were passaged every 72 hours and dissociated to smaller clumps by Gentle Cell Dissociation Reagent (Stem cell Technologies 100-0485). Upon passage, cells were seeded at 0.6 M cells/ml in mTeSR1 with 10 μM Y27632 (DNSK International, DNSK-KI-15-02).

Differentiations were started 72 hours after passage by replacing mTeSR1 media with differentiation media according to previously published protocol [1], through the end of stage 5. During feeding, cells were allowed to settle for 5 minutes, old media was aspirated away, and 300ml of warm media was added.

For plating into 24-well format, before beginning stage 5, differentiation clusters were dissociated into single cell suspension using Accutase (Innovative Cell Technologies; AT104-500), and gently disrupted by pipetting. Cells were counted and seeded at 750 K cells/cm^2^ (195K cells per well) into a Flat Bottom TC-treated 24-well plate (Falcon, 353047), coated with Matrigel (Corning® Matrigel® Growth Factor Reduced (GFR) Basement Membrane Matrix, CLS356231). Cells were fed the standard stage 5, day 1 media with the addition of 10μM Y27632 and the additional experimental chemicals. Subsequent feeds were performed with 200ul of prescribed differentiation media with the addition of tested chemicals or a blank equivalent volume of DMSO. Treatments were performed in duplicate. All experiments involving human cells were approved by the Harvard University IRB and ESCRO committees.

### RNA isolation and qPCR

RNA was extracted from entire well contents using Qiagen RNeasy kit (Cat. No. 74104), then cDNA was synthesized from 500ng of total RNA for each sample using the Invitrogen SuperScript VILO cDNA Synthesis kit (Cat. No. 11754050)

Predesigned probe/primer sets were purchased from Integrated DNA technologies. Assays were performed using TaqMan universal PCR master mix (Cat. No. 43-643-38) on a QuantStudio 7 Pro. Each cDNA was tested in technical triplicate by qPCR. Analysis was performed by calculating ΔCq values relative to UBC, then ΔΔCq values relative to DMSO control treatment, follow by fold change calculation as 2^-ΔΔCq^. Statistical analysis was performed using GraphPad Prism on fold change average values between the average of two well replicates using Dunnett’s multiple comparison test for changes from DMSO controls.

| Target transcript | Gene ID | IDT Assay ID |
| --- | --- | --- |
| Insulin | Ins | Hs.PT.58.21254175.g |
| Neuropeptide Y | NPY | Hs.PT.58.216906 |
| MafA | MAFA | Hs.PT.58.14570025.g |
| Ubiquitin C | UBC | Hs.PT.39a.22214853 |

### Bulk RNA-sequencing

Total RNA from the same extraction used for cDNA synthesis and was submitted to Admera Health (South Plainsfield, NJ), who performed rRNA depletion and library preparation with their KAPA Hyper RNA with Riboerase HMR kit. Libraries were analyzed using Illumina next generation sequencing and relative quantification was provided by Admera Health. Differential expression analysis was performed using the DeSeq2 R package, comparing each to treatment vehicle treated controls. The gseapy R package was used to evaluate enrichment of maturation neighborhood. Holm-Bonferroni method was then applied to account for the multiple testing. For projections in figure 6, the top 500 differentially upregulated genes in terms of magnitude of enrichment were projected. RNAseq results can be accessed from the NCBI GEO repository GSE309159.

## Supporting information

Descriptions of contents in supplementary data files

Lists of genes used for the generation of the maturation neighborhood in figure 2 and supplemental Figure 2.

Summary data of L1000 chemical perturbation signatures passing Structure, Quality and Overlap thresholds in Figure 3. This is the complete list of pre

Each of the 201 unique predicted compounds ranked by product value on SOM projection, compared with ranks from Clue.io and Enrichr for each compound.

Summary of compounds used in published in vitro maturation protocols and drugs annotated to have the same target. Used for generating Figure 4.

Summary of cell line data analysis. For each cell line the total number of input signatures and the number of DBSOMA generated hits from that cell lin

Summary data of L1000 genetic perturbation signatures passing Structure, Quality and Overlap thresholds for both CRISPR and Overexpression perturbatio

## Author contributions

T.R.K: Conceptualization, Methodology, Software, Formal Analysis, Investigation, Data Curation. Writing - original draft, Writing - review and editing, Visualization.

JR-F: Conceptualization, Experimental design and Resources, Writing - review and editing. Project Administration, Funding Acquisition.

## Acknowledgements

We thank M. Melé, W. Oliveros, A. Klein and M. Bulyk for their discussion and feedback on the study design, analysis approaches and the manuscript. We thank Jeffery Millman for tissue culture advice and generous donation of cells. The research was supported by an HSCI grant (DP-0215-22-00) to JR-F.

---

**Supplementary Table 1:** Predictions compared with maturation protocols.

| Protocol | Chemical | Relevant function | Compound in L100 dataset? | DBSOMA predictions | Clue.io | Enrichr |
| --- | --- | --- | --- | --- | --- | --- |
| Balboa et al [6] | ZM447439 | Aurora Kinase inhibitor | **Yes** | **ZM447439**, tozasertib | None | None |
|  | *N*-acetyl cysteine (NAC) | Antioxidant | No | cianidanol | None | None |
|  | triiodothyronine (T3) | Thyroid hormone receptor agonist | No | None | None | None |
| Liu et al [9] | ZM447439 | Aurora Kinase inhibitor | **Yes** | **ZM447439**, tozasertib | None | None |
|  | G-1 | G protein-coupled estrogen receptor | No | estrone, mestranol, tibolone, raloxifene | bazedoxifene | DPN, trilostane |
|  | H1152 | ROCK-inhibitor | No | KD-025 | None | None |
|  | CI-1033 | ERBB inhibitor | No | neratinib | neratinib, BRD-K14441456 | AG-494, tyrphostin-B44 |
|  | betacellulin (BTC) | ERBB4 agonist | No | None | None | None |
|  | 3-Deazaneplanocin A (Deza) | Histone methyltransferase inhibitor | No | None | BIX-01294 | E-7438 |
|  | ISX-9 | NEUROD1 inducer via calcium signaling stimulation | No | None | None | None |
| Rezania et al [8] | R428 | AXL inhibitor | **Yes** | **R428** | None | None |
| | Alk5iII | TGF-β type I receptor kinase (ALK5) inhibitor | No | None | SB-431542 | QX-314 |
|  | triiodothyronine (T3) | Thyroid hormone receptor agonist | No | None | None | None |

**Supplementary Table 2:** Literature search for identified candidate drugs.

| **Chemical** | **Published effect** | **Source** |
| --- | --- | --- |
| Ciclopirox | Enhances pancreatic islet health | Mihailidou et al. 2016 [34] |
| Epirizole | induced precocious beta-cell/endocrine differentiation | Rovira et al. 2011 [35] |
| Flutamide | Contributes to beta-cell proliferation | Li et al. 2008 [36] |
| Gemifloxacin | Insulinotropic potential | Ullah et al. 2022 [37] |
| Imatinib | Protects against type I diabetes.<br>Improves beta cell function | Gitelman et al. 2021 [38]<br>Xia et al. 2014 [39] |
| LY-294002 | Promotes Regeneration of Pancreatic β-cells | Xu et al. 2018 [40] |
| Milnacipran | Improves type II diabetes | Abrahamian et al. 2012 [41] |
| MK-2206 | Promotes Regeneration of Pancreatic β-cells | Xu et al. 2018 [40] |
| Moxifloxacin | Insulinotropic potential<br>Patient dysglycemia | Ullah et al. 2022 [37]<br>Chou et al. 2013 [42] |
| Neratinib | Protects beta cells from apoptosis | Ardestani et al. 2019 [43] |
| Rimonabant | Induces endocrine lineage in enteroendocrine protocol | Zeve et al. 2022 [44] |
| Roscovitine | Promotes B-cell differentiation from progenitors | Liu et al. 2018 [45] |

## Supplemental figures

**Supplemental Figure 1:**
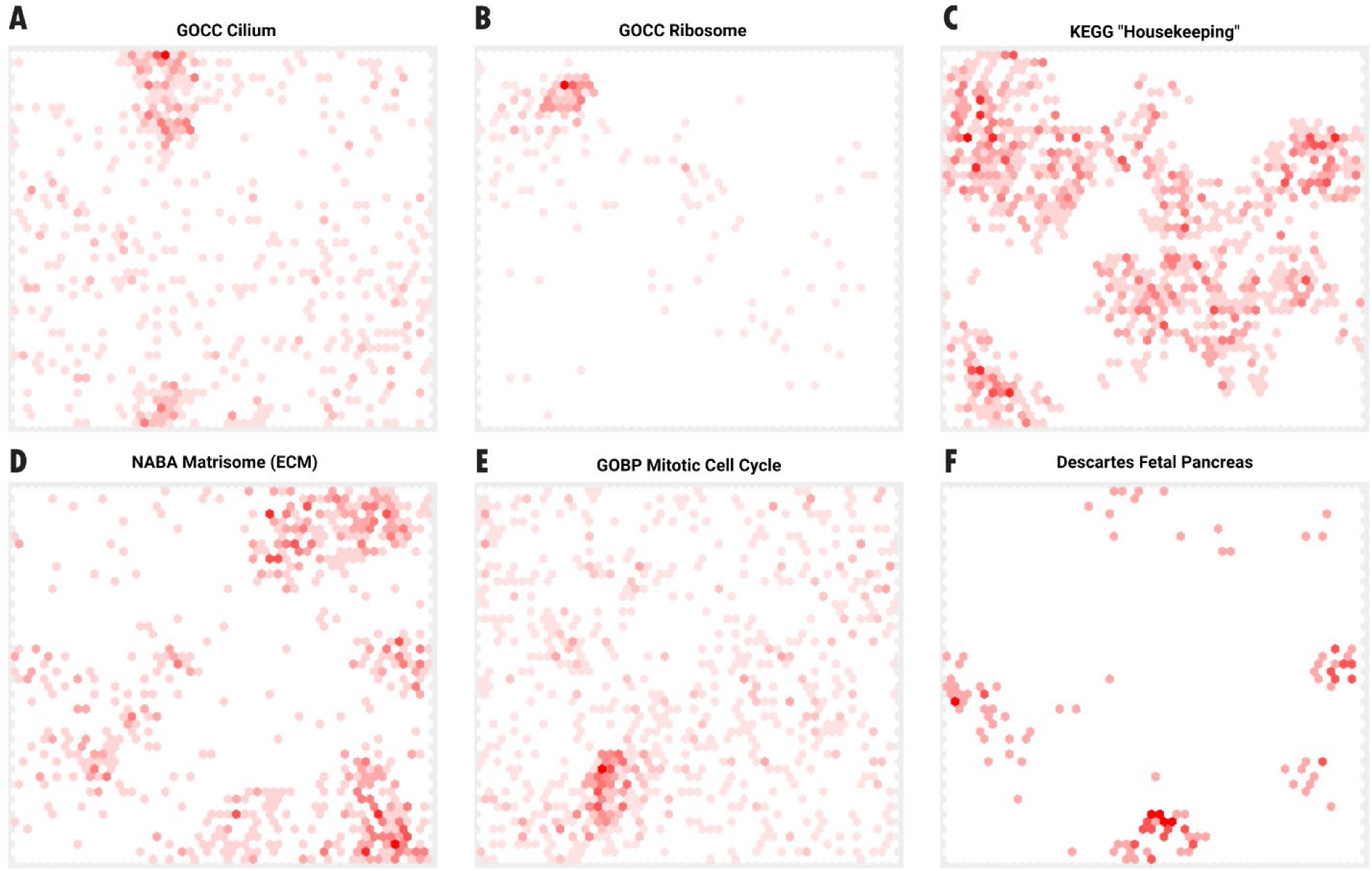
Example gene list projections on the SOM. Clustering of related genes is observed for several data sets in different regions of the map, with clusters being formed for genes related to cilium (A), ribosomal proteins (B), housekeeping (C), extracellular matrix (D) cell cycle (E) and fetal pancreas (F).

**Supplemental Figure 2:**
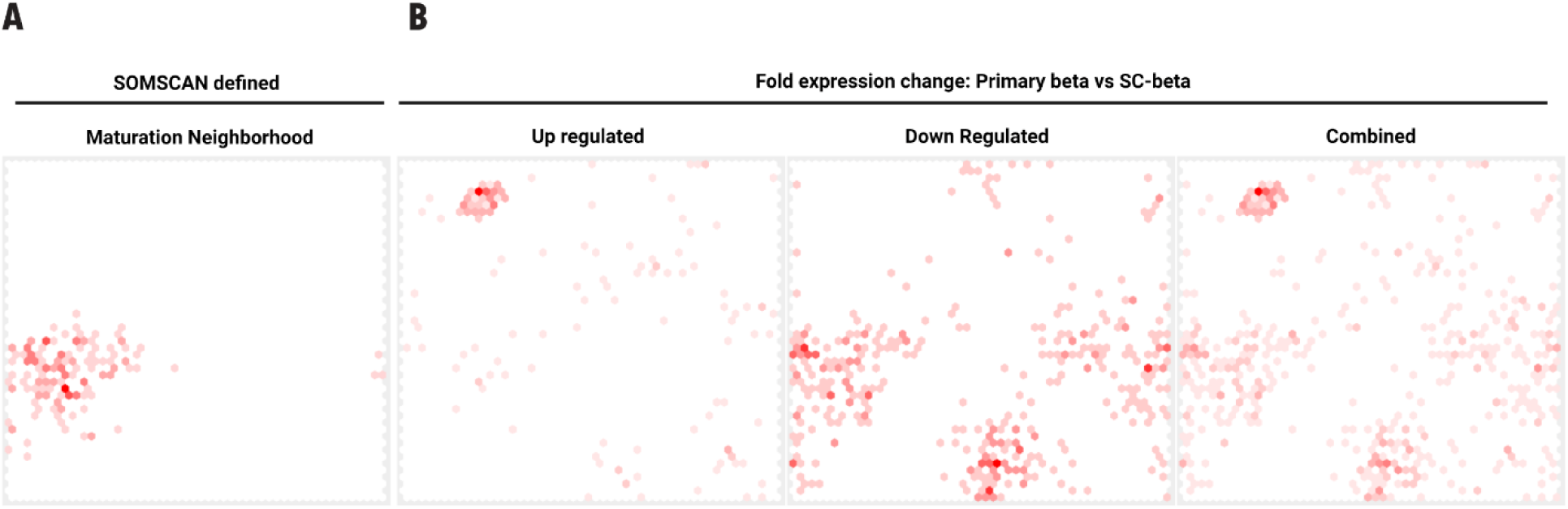
Alternative maturation gene signatures. Projection of the gene list used to define the maturation neighborhood based on SOM analysis in Figure 2 (A). An alternative approach, projecting up-regulated (left), downregulated (middle) and combined (right) gene lists of differentially expressed genes between primary β-cells and SC-β cells (B).

**Supplement Figure 3:**
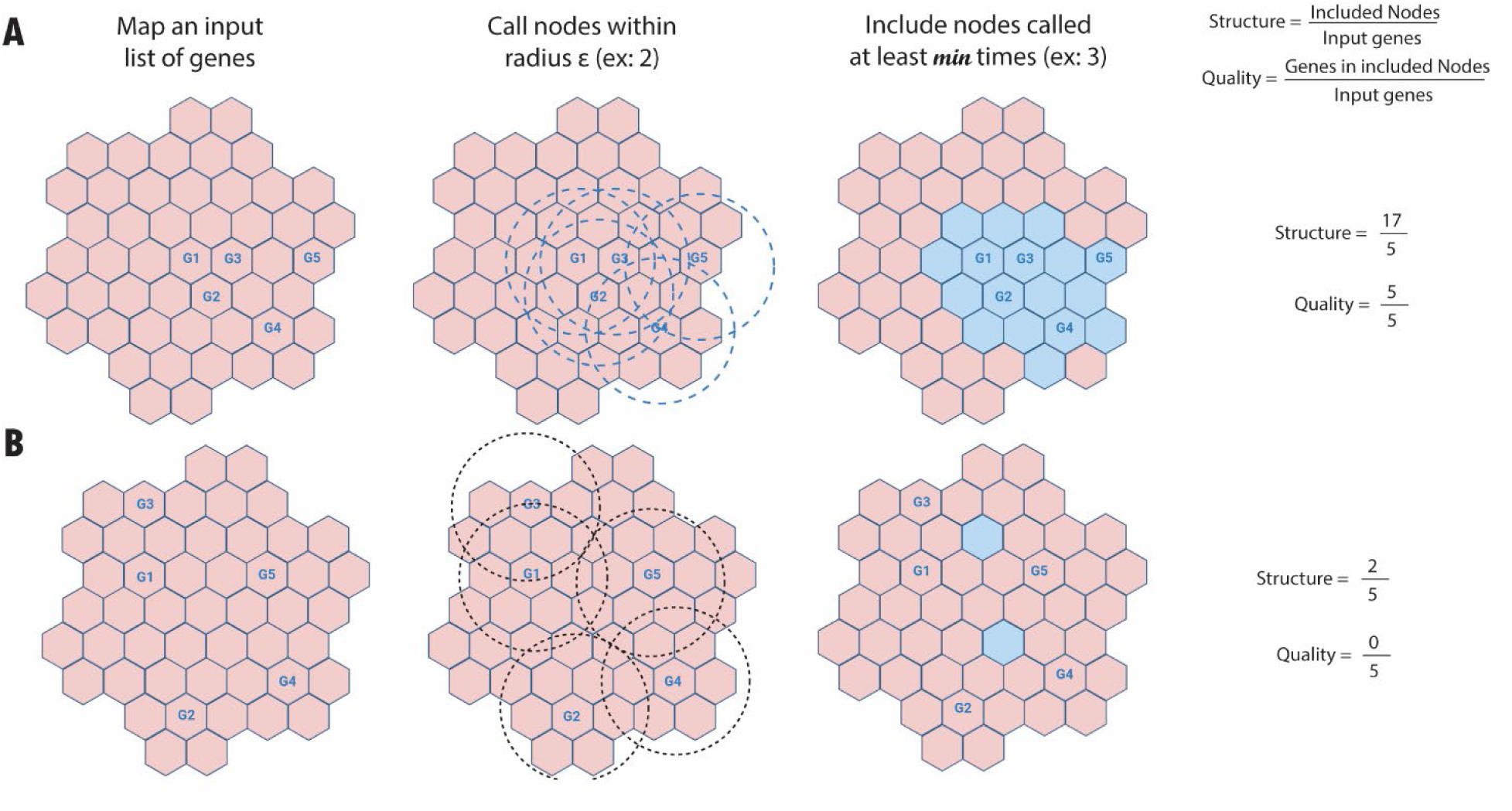
Formation of a cluster by gene lists can be scored by applying the DBSOM algorithm. DBSOM is applied here for an example space with a more clustered (A) and less clustered (B) set of inputs. DBSCAN is used to define *included* nodes (in blue). Then, the relative degree of clustering can be calculated by *structure* and the *quality* metrics, based on input genes relative to *included* nodes.

**Supplemental Figure 4:**
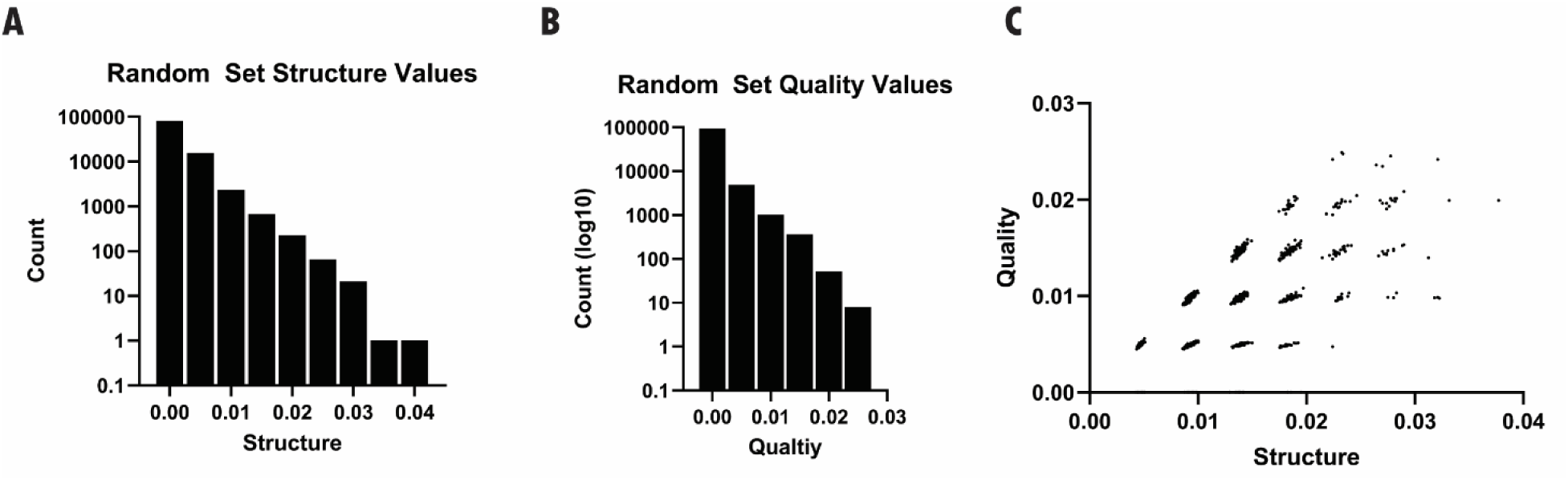
False positive estimation from random gene lists. 100,000 gene lists of size 250 were generated by random selection from SigCom dataset outputs. These sets were then evaluated for *structure* (A) and *quality* (B) metrics for comparison to real data. The values for each randomized data set are plotted together (C).

**Supplemental Figure 5:**
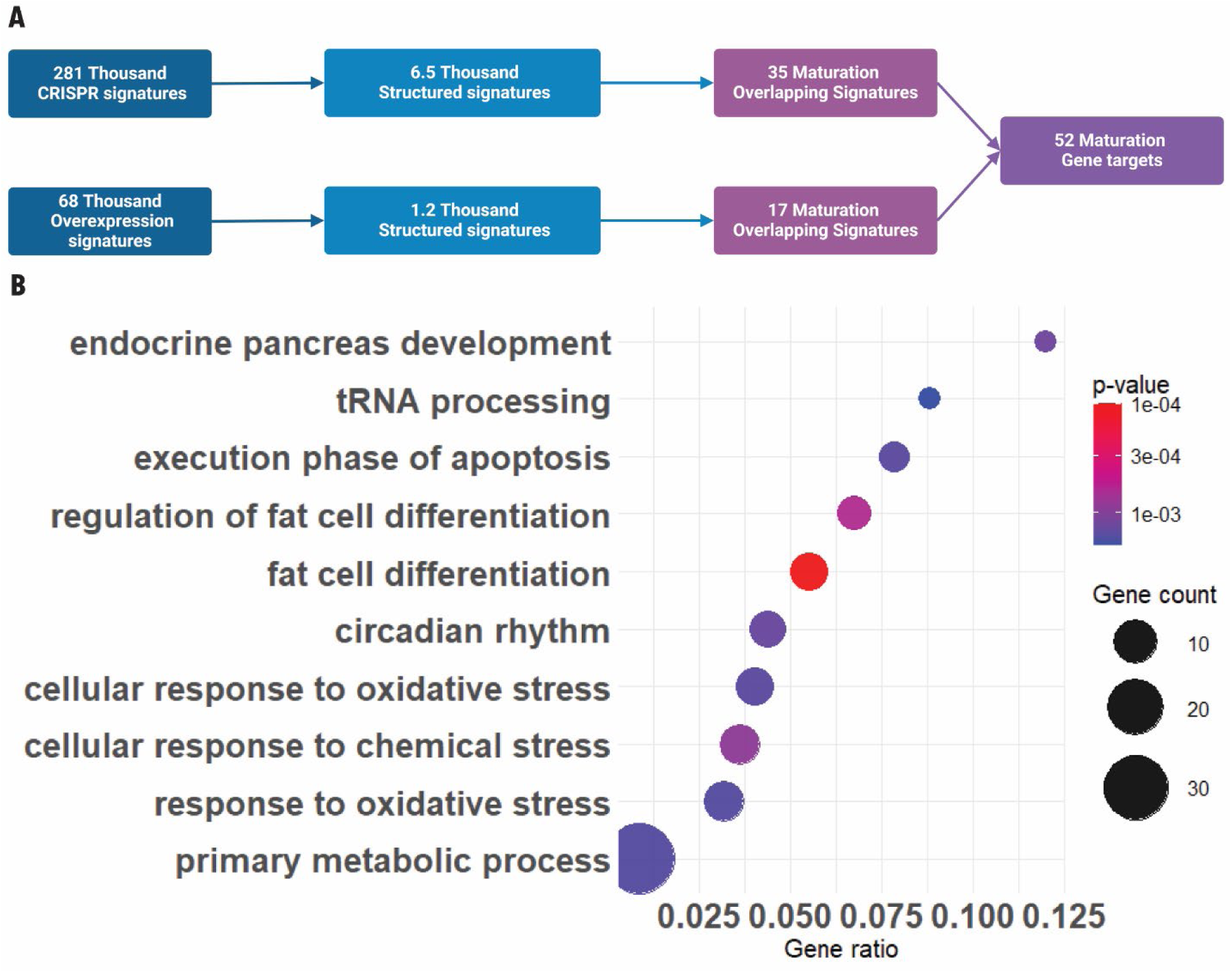
GO: enrichment analysis of predicted maturation genetic perturbations. Transcriptional response signatures for CRIPSR perturbation and overexpression perturbations were screened using the DBSOMA methodology to identify 52 putatively maturation related genes (A). Identified genes were subjected to biological process GO: term enrichment analysis, the top 16 terms, ranked by fold-enrichment, are presented (B).

