## Supplementary material for "DBSOMA: A Machine Learning Method that Identifies Chemical Modulators of Transcriptional States Uncovers Effectors of Beta-Cell Maturation": Descriptions of contents in supplementary data files

**Supplementary Data Table 1:**

Lists of genes used for the generation of the maturation neighborhood in figure 2 and supplemental Figure 2.

**Supplementary Data Table 2:**

Summary data of L1000 chemical perturbation signatures passing *Structure, Quality and Overlap* thresholds in Figure 3. This is the complete list of predicted maturation signatures. Includes information on the cell line, compound, dosage and direction of change predicted, as well as the *Structure, quality, overlap and product* calculated for each perturbation response signature.

**Supplementary Data Table 3:**

Each of the 201 unique predicted compounds ranked by *product* value on SOM projection, compared with ranks from Clue.io and Enrichr for each compound. Values of -1 indicate the compound was not found by the corresponding method.

**Supplementary Data Table 4:**

Summary of compounds used in published in vitro maturation protocols and drugs annotated to have the same target. Used for generating Figure 4.

**Supplementary Data Table 5:**

Summary of cell line data analysis. For each cell line the total number of input signatures and the number of DBSOMA generated hits from that cell line are tabulated. Used to generate Figure 3H and for the hypergeometric overrepresentation testing.

**Supplementary Data Table 6:**

Summary data of L1000 genetic perturbation signatures passing *Structure, Quality and Overlap* thresholds for both CRISPR and Overexpression perturbations. Used to generate GO:enrichment analysis presented in supplemental figure 5.

**Trained_SOM.som:**

Text file containing the trained SOM used for all analyses herein.
